# Modeling DNA opening in the eukaryotic transcription initiation complexes

**DOI:** 10.1101/2021.06.06.447228

**Authors:** Genki Shino, Shoji Takada

## Abstract

Recently, the molecular mechanisms of transcription initiation have been intensively studied. Especially, the cryo-electron microscopy revealed atomic structure details in key states in the eukaryotic transcription initiation. Yet, the dynamic processes of the promoter DNA opening in the pre-initiation complex remain obscured. In this study, based on the three cryo-electron microscopic yeast structures for the closed, open, and initially transcribing complexes, we performed multiscale molecular dynamics (MD) simulations to model structures and dynamic processes of DNA opening. Combining coarse-grained and all-atom MD simulations, we first obtained the atomic model for the DNA bubble in the open complexes. Then, in the MD simulation from the open to the initially transcribing complexes, we found a previously unidentified intermediate state which is formed by the bottleneck in the fork loop 1 of Pol II: The loop opening triggered the escape from the intermediate, serving as a gatekeeper of the promoter DNA opening. In the initially transcribing complex, the non-template DNA strand passes a groove made of the protrusion, the lobe, and the fork of Rpb2 subunit of Pol II, in which several positively charged and highly conserved residues exhibit key interactions to the non-template DNA strand.

**Author Summary:** Transcription is fundamental phenomenon in all species, and its regulation of the initiation process is important. In eukaryotes, multiple proteins, which is not only RNA polymerase but also transcription factors, assemble on the promotor to form the complex. After that, this complex open the part of DNA duplex, leading to forming single-stranded region for transcription. Previous study, including structural analysis and biochemical experiments, obtained the information about single-stranded region, but the structure is incomplete, so the details of the mechanism remains unclear. In this study, by the calculation using structural information about the protein-DNA complex, we obtained the structure of the complex with complete single-stranded region. As the result of the calculation, it found that the size of single-stranded region was fluctuated, which is related the sequence of the region. Furthermore, we found the intermediate state during the extended process of single-stranded region, which has not been reported in previous study. In this state, we found that the particular region of RNA polymerase is relevant to the extension of single-stranded region. These founding may contribute the understanding about the regulation of transcription initiation.

## Introduction

Transcription is fundamental to virtually all area of biology. In eukaryotic cells, RNA polymerase II (Pol II) transcribes all messenger RNAs, making it of central importance. The Pol II transcription initiation requires progressive assembly of several general transcription factors (TFs) and Pol II on the promotor DNA sequence, forming the pre-initiation complex (PIC). After the initial transcription of short RNAs, the transcription machinery escapes the promoter region converting its architecture for the transcription elongation. Much of the transcriptional regulations are related to these early stages of transcription and thus it is of utmost importance to understand the molecular mechanisms of the transcription initiation, which we focus in this study.

Overall processes in the Pol II transcription initiation have been characterized via decades of studies. The PIC consists of Pol II and six general TFs, TFIIA, TFIIB, TFIID, TFIIE, TFIIF, and TFIIH [1, 2, 3, 4]. In addition, coactivators such as Mediator are involved in its regulation [5, 6]. The initiation process begins with the recognition of the promoter DNA sequence by TFIID. For the promoter sequences that contain the classic TATA box, the TATA-binding protein (TBP) in TFIID binds the TATA box DNA sequence, leading to ~90-degree bend of the DNA. Then, TFIIA, TFIIB, Pol II-TFIIF complex assemble in this bent site. Further, TFIIE and TFIIH are recruited in order, to form the PIC with the bent duplex DNA (termed the closed complex, CC). In particular, PIC without TFIIH is called as core PIC (cPIC). Next, the promoter DNA melts into the template and non-template DNA strands, driven by the ATP-dependent translocase activity of TFIIH (termed the open complex, OC). The template DNA strand moves toward the active site of Pol II. The melted DNA region is called “DNA bubble”, of which size is experimentally characterized as ~6 bp in the OC state [7]. Subsequently, the DNA bubble expands to ~13 bp [7], which allows the template DNA strand reaching to the active site to begin the messenger RNA synthesis. The complex in which the initial transcription begins is called the initially transcribing complex (ITC). Notably, while the ATP-dependent translocase activity of TFIIH facilitates the promoter DNA opening [8, 9], some promoter DNAs can open spontaneously without TFIIH [10].

Recently, the cryo-electron microscopy (cryo-EM) revealed near-atomic structures in key stages of the Pol II transcription initiation [5, 6, 10], which provided the model of DNA opening process in the PICs (Fig. 1A) [10]. The model is based on the yeast CC, OC [10], and ITC structures [11], and the highly conserved human CC structure [12]. However, the state transitions from CC to OC, and to ITC were not directly observed. Moreover, the modeled structures of OC and ITC by cryo-EM do not contain parts of DNA strands because of high flexibility in the DNA bubble. Thus, how the template and non-template DNA strands behave inside Pol II has not been fully understood. Complementarily, the DNA bubble size in OC and ITC states has been detected via optical and magnetic tweezer experiments [7, 13]. However, the structural details in the DNA bubble is currently missing.

**Figure 1:**
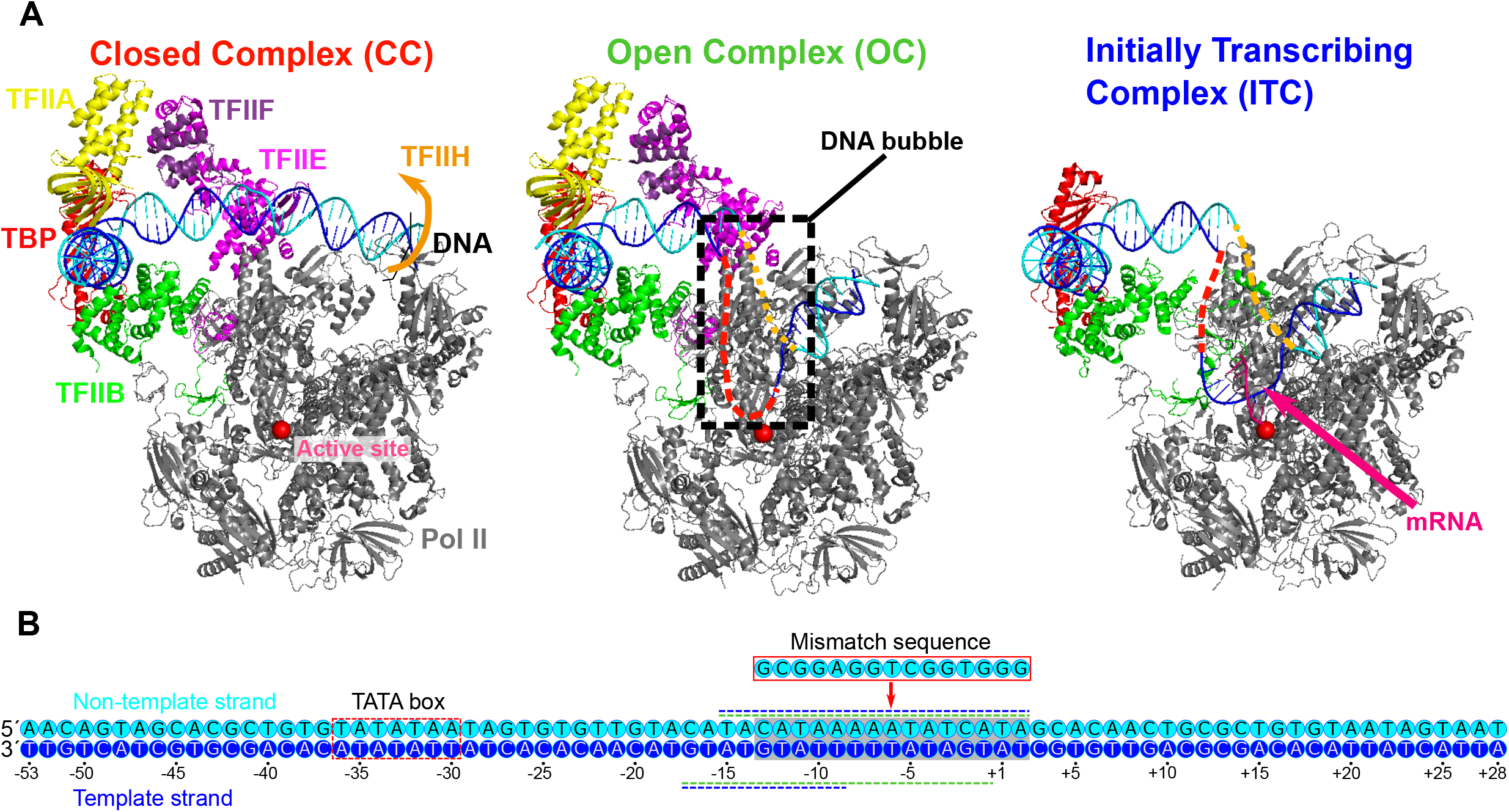
Three yeast PICs of RNA polymerase II and the promoter sequence used. (A) The three PICs obtained by cryo-EM; the closed complex (CC) (PDB: 5FZ5) (left), the open complex (OC) (PDB: 5FYW) (center), and the initially transcribing complex (ITC) (PDB: 4V1N) (right). The CC and OC models contain the promoter DNA, Pol II, TBP, TFIIA, TFIIB, TFIIE, and TFIIF. The ITC model contains the promoter DNA, Pol II, TBP, TFIIB, and 6 bp nascent RNA. Parts of the melted DNA were not modeled in the OC and ITC states (Red and orange broken lines). (B) The promoter DNA sequence used in the current study (numbered relative to the transcription starting site). The sequences are taken from those used in the cryo-EM studies of the OC and the ITC [10, 11]. Blue, the template DNA strand; cyan, the non-template DNA strand; red dashed square; TATA box; gray square, region to which mismatched sequence is introduced in a simulation; green and blue horizontal dashed lines along the sequence, the regions not appeared in the OC and ITC models by cryo-EM, respectively.

Given such situations, molecular dynamics (MD) simulations can offer another complementary approach to address the structural dynamics of the Pol II transcription initiation since MD simulations can provide high-resolution spatiotemporal information [14, 15, 16, 17, 18]. However, since the DNA opening process involves rather large-scale and slow movements of DNA within large complexes, conventional MD simulations with fully-atomic resolution (designated as the all-atom (AA) MD hereafter) cannot easily sample these structural dynamics. To circumvent this difficulty, one can alternatively use coarse-grained (CG) MD simulations, which can speed up the simulation by orders of magnitude at the cost of accuracy. Once comprehensively sampled by CG-MD, one can back-map the sampled CG structure models into AA models, followed by AA-MD simulations. A recent study employed such a protocol to gain comprehensive and high-resolution energy landscape in a bacterial RNA polymerase [19].

In this study, using the cryo-EM yeast structures for the CC, OC, and ITC, we performed multiscale MD simulations to model structures and dynamic processes of DNA opening. Combining CG- and AA-MD simulations, we first obtained the atomic model for the DNA bubble in the OC. Then, in the CG-MD simulation from the OC to the ITC, we found a previously unidentified intermediate state which is formed by the bottleneck in the fork loop 1 of Pol II: The loop opening triggered the escape from the intermediate, serving as a gatekeeper of the promoter DNA opening. In the ITC, the non-template DNA strand passes a groove made of the protrusion, the lobe, and the fork of Rpb2 subunit of Pol II, in which several positively charged and highly conserved residues exhibit key interaction to the non-template DNA strand.

## Results & Discussions

### Multiscale modeling of pre-initiation complexes

Our multiscale modeling begins with CG-MD simulations that connect the three states of the PIC; the CC, OC, and ITC. The constructed CG models were then back-mapped to AA models, which is followed by short-time MD simulations with the AA models.

We employ the CG model that has been extensively used to protein-DNA complexes [20, 21, 22, 23, 24, 25, 26, 27, 28]. In the CG model, each amino acid in proteins is represented as one particle located at the Cα atom position and each nucleotide in DNA is modeled by three particles each representing the phosphate, the sugar, and the base. The protein energy function AICG2+ contains the contact potentials that stabilizes the predefined reference (native) structure, i.e., the structure-based model [29]. The DNA energy function 3SPN.2 is empirically tuned to reproduce several experimental data such as the sequence-dependent melting temperature and bending modulus [30]. The protein-DNA energy function consists of the generic terms; the short-range repulsion and the electrostatic interaction, and the specific interactions; structure-based contact potentials (See Materials and Methods for more details).

The simulation system consists of an 81-bp promoter DNA (the sequence shown in Fig. 1B) and the protein complex that contains Pol II, TBP, TFIIA, TFIIB, TFIIE, and TFIIF (that is, this study deals with cPIC). TFIIH is not included because the DNA bubble can form without TFIIH [10, 31] and because the structural information on ATP-dependent conformational change in TFIIH is incomplete albeit some structures previously reported [5, 32, 33]. The root-mean-square-differences (RMSD) were 0.85Å between CC and OC, and 3.5Å between OC and ITC, which are smaller than the resolutions reported in the cryo-EM analysis (8.8 Å, 4.4 Å, and 7.8 Å, for CC, OC, and ITC models, respectively). In the CG-MD simulations, these modest-sized structure changes in proteins should appear via the interaction to DNA (and a short RNA in the case of ITC). Since CC contains the weakest protein-DNA interaction among the three complexes structure models, we took the protein structure of CC as a reference structure of the protein complex in the CG model throughout this study. For the specific protein-DNA interaction, we collected protein-DNA contacts in the three complexes structures and used its union set for the structure-based contact potential.

### Modeling the DNA bubble in the open complex

First, to obtain the OC model with the open DNA, we performed 40 independent CG-MD simulations of 5 × 10^6^ MD steps, starting from the CC structure (Fig. S1). In any of the simulations, the promotor DNA did not melt spontaneously and most of the OC specific protein-DNA contact did not appear although the particular region of the promotor (−18~ +7 relative to the transcription start site (TSS)) was distorted toward the cleft of Pol II (Fig. S1). This distortion in DNA was observed in a previous cryo-EM study, which indicates the pre-stage of DNA opening [32]. A previous study shows that the DNA opening in the absence of TFIIH takes a very long time; the real-time observation of the formation of the DNA bubble shows that it takes a few seconds [13]. Therefore, it is reasonable that we did not observe spontaneous DNA opening in our CG-MD simulations that cannot cover second time scales.

Then, to enforce the prompt DNA opening, we modified the non-template DNA sequence to introduce the DNA mismatch of 15 bases into the promotor (Fig. 1B). The introduced mismatch is identical to that used for the cryo-EM structures of OC and ITC [10, 11]. Under this condition, we performed 40-independent CG-MD simulations of 2 × 10^7^ MD steps (Fig. 2). Figure 2B left panel depicts a representative time course of the fraction of protein-DNA contacts specific to CC (red) and to OC (green). In this trajectory, in the very initial phase, ~80% of the CC-specific contacts were lost, whereas ~40% of the OC-specific contacts were formed to reach an intermediate state, which we call “pre-OC” state (Fig. 2A center). In the pre-OC state, most of the mismatched DNA region melted to form a bubble of ~13 bp (Fig. 2C). This caused the +2 site of the template DNA strand to form new contacts with Pol II (see Movie S1). All the 40 trajectories paused at this pre-OC state (40 / 40 cases). In the trajectory in Fig. 2B left panel, the template DNA strand jumped further down to the active site at ~ 0.9 × 10^6^ MD steps, reaching to the OC-like state with the mismatch (Fig. 2A right, Fig. 2B) (22 / 40 cases. Of the 22, we observed that the complex further moved towards the ITC state in 5 cases). The transition was driven by the new contact formation of the +1 site of the template DNA strand with Pol II (see Movie S1).

**Figure 2:**
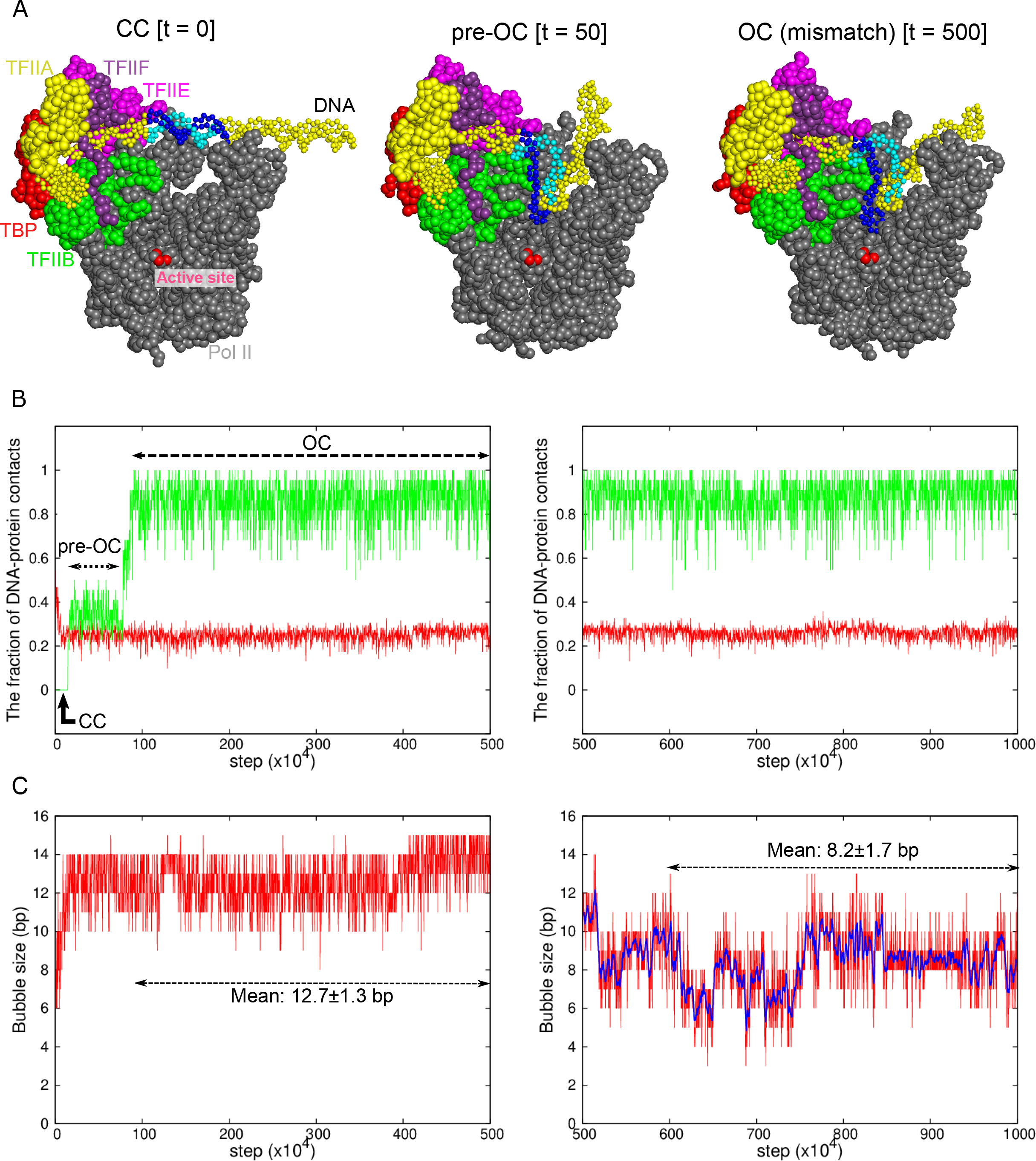
Coarse-grained MD simulation for the transition from the CC to OC states. Results of a representative trajectory are shown. (A) Snapshots at t = 0 (left, the CC state), at t= 50 × 10^4^ MD steps (center, the pre-OC state), and at t = 500 × 10^4^ MD steps (right, the OC state with the mismatch). Some proteins are not displayed to make DNA visible. The same colors are used as Fig. 1A. Blue and cyan region in DNA indicates the 15-bp mismatch region, forming the DNA bubble. (B) The time course of the fractions of protein-DNA contacts specific to CC (red) and OC (green). (C) The time course of the DNA bubble size. In (B) and (C), the left (right) panels are from the first (second) halves of MD simulations with (without) the DNA mismatch. The blue curve in the right panel in (C) shows a moving average over 11 points.

To obtain the OC structure model without the DNA mismatch, for an obtained OC-like structure with the mismatch, we changed the DNA sequence back to the original sequence without mismatch, followed by 5 × 10^6^ MD step CG-MD simulations. A representative trajectory is depicted in the right panels of Fig. 2BC. While the overall positioning of DNA did not change (Fig. 2B right), part of the melted DNA regained the base pairing during the trajectory (Fig. 2C right). The observed bubble size fluctuated in time in the range of 6-10 bp, with the mean and the standard deviation 8.2±1.7 bp. Notably, we observed that the bubble size depends on the promoter sequence to some extent; with the promoter sequence used in the single-molecule magnetic tweezer experiment, our simulation resulted in the bubble size of 5.5 ± 1.4 bp, which is fairly compared with the experimental estimate, 6.1 ± 0.3 bp [7].

To detect protein-DNA interactions in the OC state at atomic level, we modeled all-atom structures via back-mapping from the snapshots of a CG-MD trajectory with the DNA bubble sizes of 6 bp and 9 bp. The obtained all-atom models were further relaxed/refined by 10 ns MD simulations with explicit water solvent (Fig. 3). In the upstream of the DNA bubble, we find that the hydrogen bonds of the TFIIE E-wing residues K80 with the non-template DNA at −11 to −10 sites, and with the template DNA at −13 site, which are present in the CC state, are maintained (Fig. 3B). These interactions are suggested to facilitate the promoter opening and contribute to the efficiency of transcription initiation [34]. Comparing the structures with 6 bp and 9 bp DNA bubbles, we find that the template DNA strand is rather similar each other while the non-template DNA strand is more mobile. The 6 bp in the downstream side (from −4 to +2 sites) were melted in both structures, while the 3 bp in the upstream side (from −7 to −5 sites) were formed/melted in the 6 bp/9 bp DNA bubbles.

**Figure 3:**
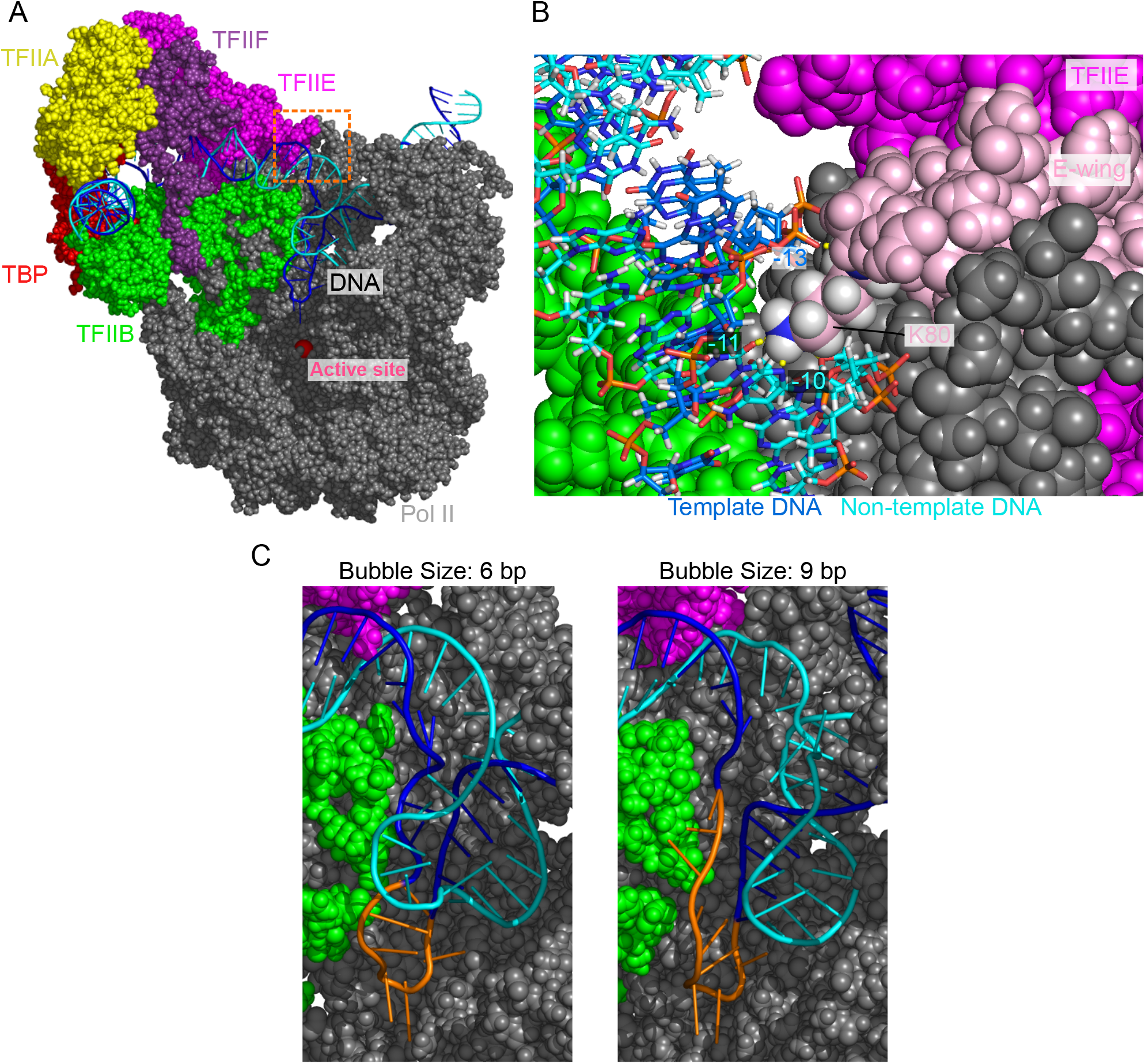
The open DNA in the OC state of PIC. (A) Atomic structure model for the OC state. Some proteins are not displayed to make the DNA visible. (B) A close-up view of the orange dashed squared area in (A). Pink, the E-wing of TFIIE; green dashed circle, hydrogen bonds between DNA and the E-wing. (C) Open DNA structures with the bubble size of 6 bp (left) and 9 bp (right). Orange, the template DNA strand in the bubble.

### Dynamical modeling of the transition from the open complex to the initially transcribing complex

Next, we addressed dynamic process of the transition from the OC state to the ITC state. Starting from a final snapshot of the previous simulation that paused at the OC state for the promoter DNA without the DNA mismatch, we performed 140 independent CG-MD simulations of 2 × 10^7^ steps (Fig. 4). In most trajectories (130 /140 cases), the contact between the promoter DNA (from −16 to −9 sites) and the E-wing of TFIIE persisted for the whole simulation time, which clearly precluded the template DNA strand from accessing the active site. Only in 10 cases, we observed the disruption of this contact, which directly triggered the template DNA strand to move down towards the active site (Fig. 4). Nine out of these 10 trajectories reached the ITC state.

**Figure 4:**
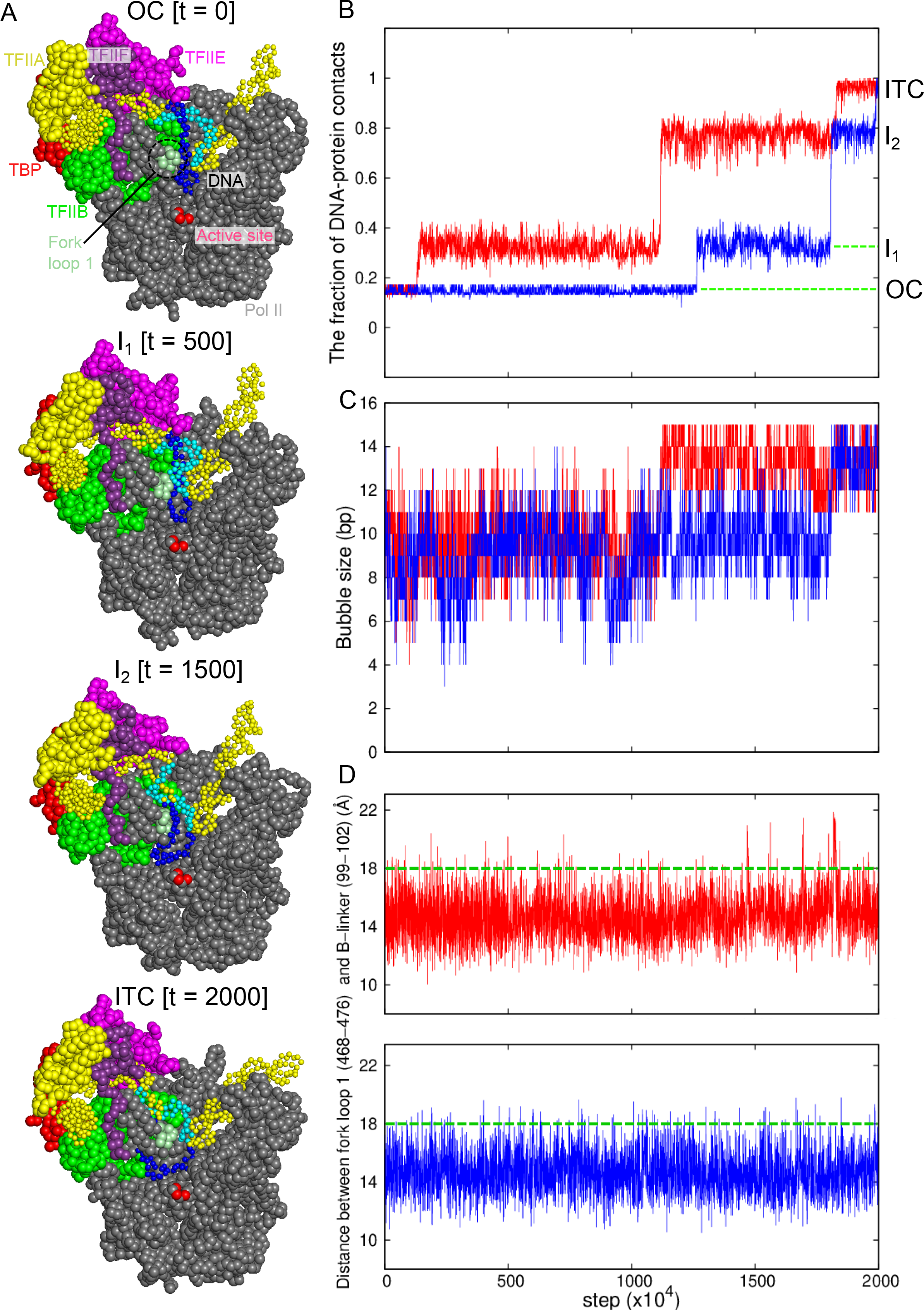
Coarse-grained MD simulation for the transition from the OC to ITC states. Results of two representative trajectories are shown in red and blue curves. (A) Snapshots from the red trajectory at t=0 (top, the OC state), at t=500 × 10^4^ MD steps (I_1_ state), at t=1500 × 10^4^ MD steps (the I_2_ state), and t=2000 (bottom, the ITC state). (B) The time course of the fractions of protein-DNA contacts specific to ITC. (C) The time course of the DNA bubble size. (D) The time courses of the distance between the centers of mass of the fork loop 1 of the Pol II Rpb2 (468-476 residues) and the B-linker in TFIIB (99-102 residues). Green dashed lines, a characteristic distance for the template DNA to pass through the fork loop 1.

In these successful trajectories, we found two intermediate states (Fig. 4A the second and the third models) before reaching the ITC state (Fig. 4A bottom). In a representative trajectory (red in Fig. 4B-D), the first transition occurred at ~1 × 10^6^ MD steps, after which about 35% of the ITC specific contacts were formed. In this intermediate state I_1_, ~4 bp of the template DNA strand, (−2 ~ +2 sites, relative to the TSS) approached the active site, while the contact between the DNA and the TFIIE E-wing is maintained (Fig. 4A the second structure). After a long waiting time, the DNA was detached from the TFIIE E-wing region (at 1.1 × 10^7^ MD steps in the red trajectory), followed by the motion of the entire DNA bubble towards the active site (Fig. 4A the third structure). However, the template strand DNA in the upstream side of the DNA bubble, −13 ~ −9 sites, collides with the fork loop 1 of Rpb2 subunit of Pol II (Fig. 4A, the third structure, and Fig. S3 left). This forms a metastable intermediate state I_2_. After some duration time at this intermediate state, the complex made the final transition to the ITC state (at ~ 1.7 × 10^7^ MD steps in the red trajectory). The other successful trajectories followed similar pathways.

To increase the samples of transitions to the ITC state, we performed 160 extra-simulations of 5 × 10^6^ MD steps in which the contact between DNA and the TFIIE Ewing was weakened intentionally (see Materials and Methods, Fig. S2). In this setup, we observed the successful transition to the ITC state for 102 out of 160 cases with the transition pathway unchanged. The rest of trajectories stayed at the intermediate state I_2_ until the end of trajectories (58 / 160 cases).

In the ITC state, the DNA bubble size was, on average, 13.4±1.1 bp (Fig. 4C & Fig. S2B), which perfectly agrees with the previous estimate (13. 4 bp) [7].

### Fork loop 1 serve as a gatekeeper

The intermediate state I_2_ appears because of the blockage by the fork loop 1, which led us to hypothesize that the fork loop 1 may serve as a gatekeeper. To monitor large-scale motions of the fork loop 1, we plotted in Fig. 4D the time courses of the distance between the fork loop 1 and the B-linker of TFIIB, finding that the fork loop 1 exhibits intermittent large-scale fluctuation to open the gate (green dashed lines in Fig. 4D). In the representative time course (red), the time of the transition from I_2_ to ITC states in Fig. 4B coincides with a large-scale opening. Looking into structure changes at the time, we found that the template DNA strand passed the fork loop 1 upon the loop opening, and moved toward the active site (Fig. 4A, Fig. S3 right). Notably, in any trajectory, the non-template DNA strand never passed the fork loop 1. Instead, the non-template DNA strand approached to the wall of Pol II. Therefore, after the passage of the template strand, the fork loop 1 is located inside the DNA bubble. This support the hypothesis that the fork loop 1 serve as a gatekeeper; it is only passed by the template, but not the non-template DNA strand. This role is supported by previous studies [10, 35, 36].

The fork loop 1 sequence is fairly well conserved from yeast to human (Fig. 5). In the human Pol II, it has been reported that a mutant that deletes two residues in the fork loop 1 (K458, A459 in human Pol II, which align with K471, A472 in yeast Pol II) abolishes the transcription in vitro [37]. This supports the crucial role of the fork loop 1. The mutation may alter the loop opening dynamics, which led to the malfunction of Pol II.

**Figure 5:**
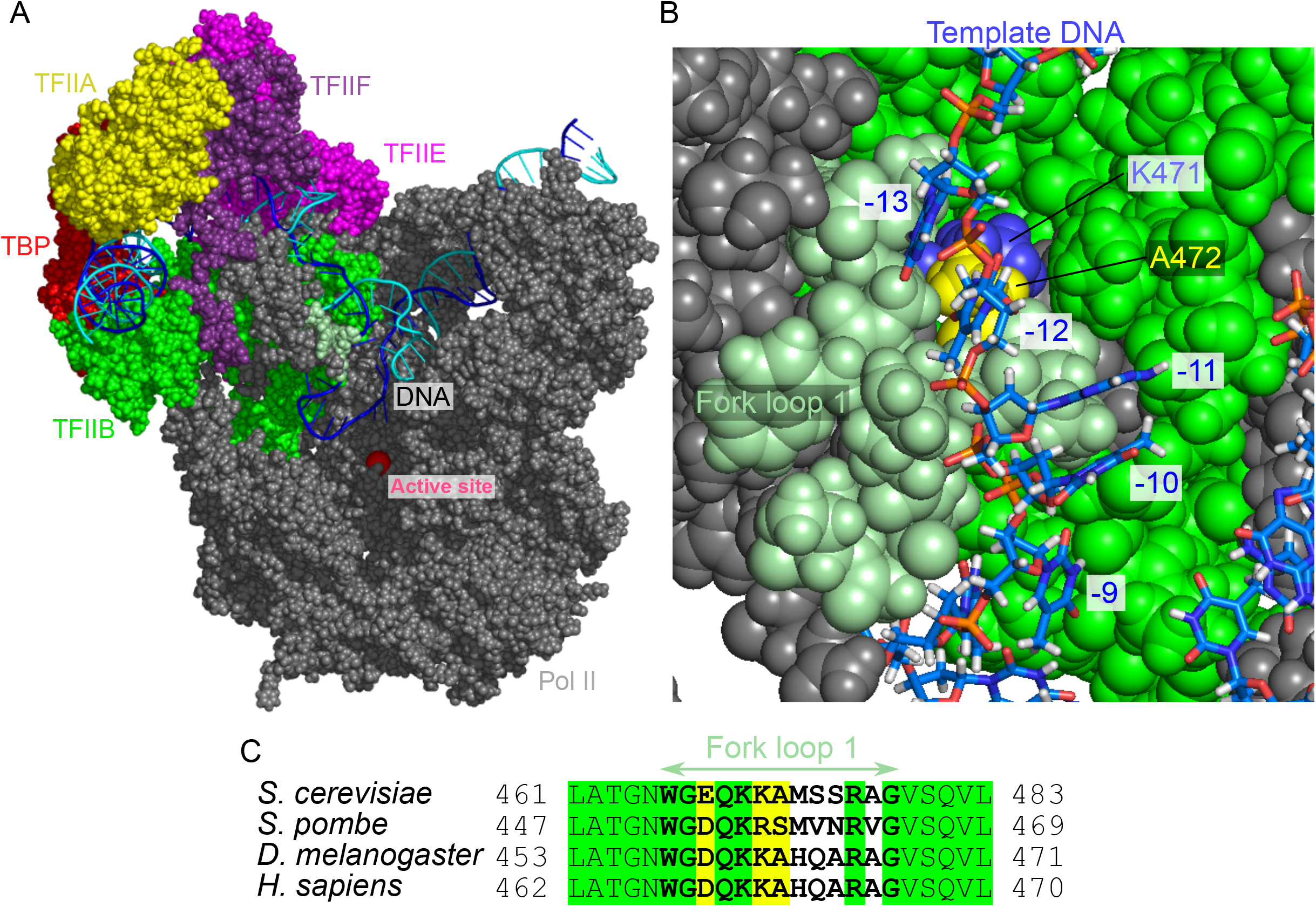
The open DNA in the intermediate state I_2_. (A) The atomic structure model for the I_2_ state. Some proteins are not displayed to make the DNA visible. (B) The close-up view of the fork loop 1 (pale green) that blocks the template DNA passage. (C) Multiple sequence alignment of the fork loop 1 region of the Rpb2. Green, invariant; yellow, conserved.

### The template and non-template DNA strands in the initially transcribing complex

To predict the placement of the template and non-template DNA strands and probe protein-DNA interactions in the ITC state, we constructed all-atom structure model via the back-mapping from snapshots in a CG-MD trajectory (Fig. 6A), which is followed by 10 ns AA-MD simulations. We note that the template DNA strand was anticipated to be in the wall from the cryo-EM study even though the cryo-EM structure model for the ITC state does not contain the segment of the template and non-template DNA strands [10]. The constructed model in this work supports this prediction; the template DNA strand is indeed placed in the wall of Pol II (Fig. S3 right). More specifically, our model suggests that the non-template DNA strand is localized at the protrusion, the lobe, and the fork of RPB2 subunit of Pol II (Fig. 6BC). This placement of the non-template DNA strand is fairly close to that found in the yeast elongation complex structures solved by X-ray diffraction (Fig. S4) [38]. These regions form the groove with many basic amino acids (Fig. 6C). Along the groove the three basic residues, R241, R504, and K507 made specific interactions to the DNA at −1, +1, and +2 sites (Fig. 6C). These three residues are strictly conserved across broad range of eukaryotes (Fig. 6D).

**Figure 6:**
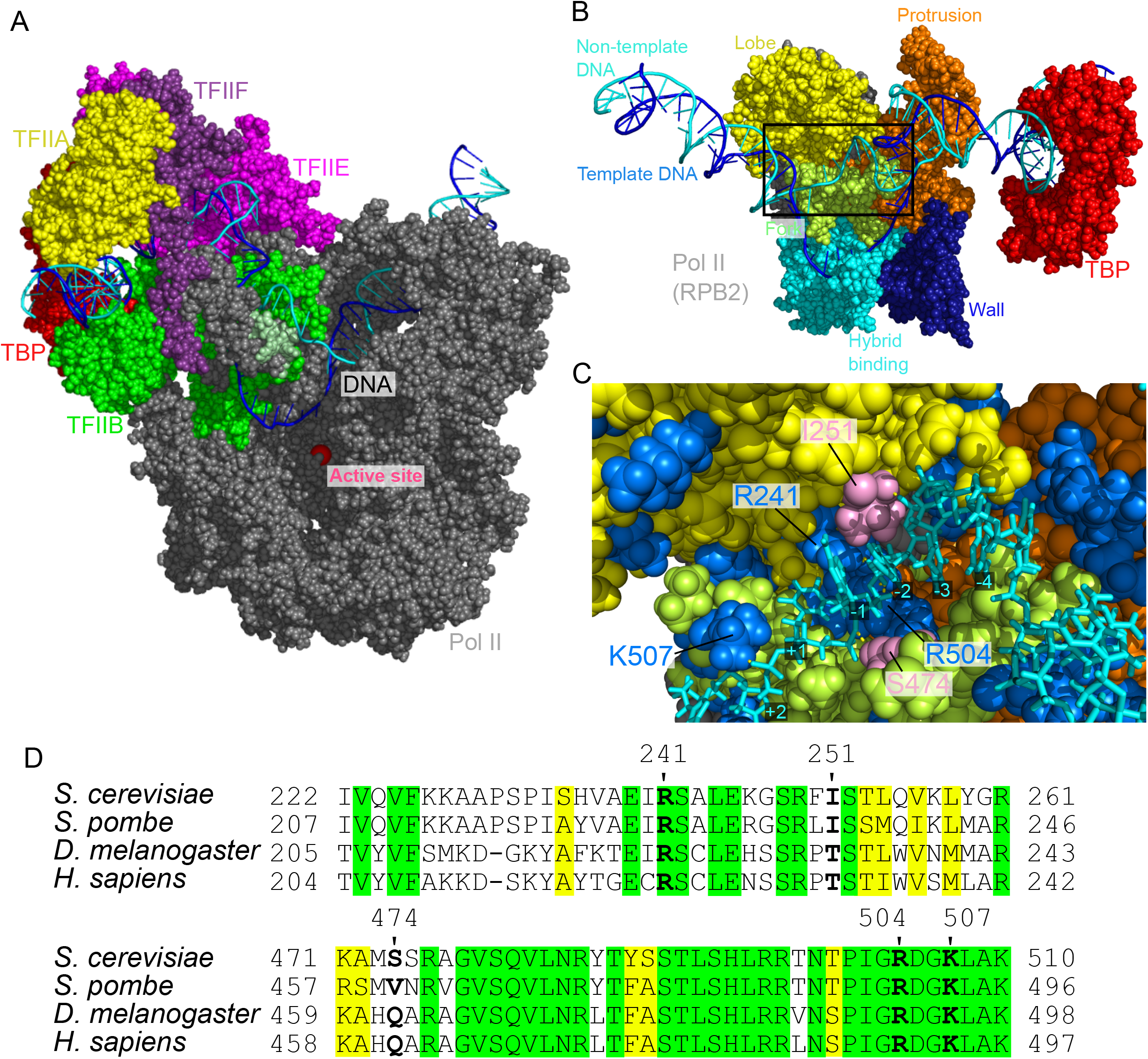
The open DNA in the ITC state of PIC. (A) The atomic structure model for the ITC state. Some proteins are not displayed to make DNA visible. (B) The atomic structure model from the back side of (A), which focuses the non-template DNA strand path. (C) The close-up view of the Rpb2 and non-template DNA strand in the squared area in (B). Blue, positively charged residues; pink, I251 and S474 that form hydrogen bonds to DNA; yellow dashed lines, hydrogen bonds between bases of the non-template DNA strand and amino acids. (D) Multiple sequence alignment of the residues around those shown in (C). Green, invariant; yellow, conserved.

### Biological Insight

Our computational modeling revealed that, after passing the OC state, the PIC passes two intermediate states before reaching the ITC, through which a small DNA bubble in the OC is expanded to complete the DNA bubble ready for RNA synthesis. One key gating state is I_2_, where the upstream part of the template DNA strand (−9 to −13 sites) interacts with the fork loop 1. The fluctuation of fork loop 1 was obligatory to escape from I_2_ that leads to engaging the template DNA into the active center at ITC (Fig. 4, 5). The non-template DNA strand did not pass the fork loop 1, suggesting that the fork loop 1 serves as the gatekeeper for the DNA bubble.

Previous studies implicated two critical roles of fork loop 1. Based on the structural change in the fork loop 1 between the nucleic-acid free state and in the transcribing state [35, 36] as well as the mutation assays (cite), the fork loop 1 was considered to play an important role in the transcription initiation. Alternatively, since the fork loop 1 is located at around the terminus of the DNA/RNA hybrid in the elongation complex, it may have important roles in separation of the product RNA from the template DNA strand. Our current simulation clearly supports the former functional role. The fork loop 1 forms a gate together with the rudder of RPB1 subunit in Pol II and serves as a gatekeeper for the engagement of the template DNA strand, but not the non-template DNA strand. The structural model obtained can be used to guess key residues as the gatekeeper, which can be examined by mutagenesis experiments.

Moreover, the current study found that non-template DNA strand in ITC is localized in the groove, formed by the protrusion, the lobe, and the fork of RPB2 subunit (Fig. 6). This pass is close to the non-template DNA strand pass in the yeast elongation complex [38], suggesting its ubiquitous importance. However, to our knowledge, these interaction sites were not investigated by mutagenesis. Systematic mutation assays in these sites would clarify the roles of stabilizing the non-template DNA strand in the transcription process.

### Conclusion

By multiscale molecular modeling and MD simulations based on cryo-EM structure models, we modeled DNA opening in the yeast PICs. We first obtained an atomic model for the DNA bubble in the OC, which exhibited the bubble size consistent with the previous experimental estimate. In the dynamic simulation from the OC to the ITC, we found an intermediate state in which the fork loop 1 blocks the template DNA strand passing into the active site. We found that, in the constructed model of ITC, the non-template DNA strand fits into the groove made of the Rpb2 of Pol II.

Obviously missing in the current work is the kinetic and energetic arguments on the very initial process of the DNA opening in the transition from the CC to the OC states. In this study, even with the use of CG-MD simulations, the DNA opening was too slow to be simulated directly in MD simulations for the native promoter sequence. Instead, we needed to introduce a mismatch sequence in the promoter region. This is clearly a limitation. To study kinetic and energetic aspects in this initial DNA opening without the mismatch sequence, we need some advanced sampling methods, such as the umbrella sampling, the Markov-state modeling [39], and the string method [40]. Alternatively, since the ATP-driven motor activity of TFIIH helicase is expected to accelerate the DNA opening, including this effect either explicitly or implicitly may enable to simulate the dynamic process of the bubble formation more directly. These developments are left for future studies.

Related to this, it has been known that Pol I and Pol III systems do not have TFIIH-like helicases [41, 42, 43, 44], yet initiating the transcription efficiently via similar three states [45, 46, 47]. Comparison of the transcription initiation in the three RNA polymerase systems can put forward comprehensive understanding of transcription initiation in Eukaryote.

## Materials and Methods

### Preparation of the simulation system

We modeled the three yeast structures, CC, OC, and ITC, based on the cryo-EM structure models, 5FZ5 [10] and 6GYL [32] for CC, 5FYW [10] for OC, and 4V1N [11] for ITC. Missing residues in the original models were modeled by the software MODELLER [48, 49, 50, 51].

We used the DNA sequence identical to that used in cryo-EM studies [10, 11]. The sequence was derived from the promoter sequence of HIS4 gene locus, from which 28 bp were deleted at the downstream of the TATA box.

### Coarse-grained MD simulations

In this study, we applied the coarse-grained (CG) simulation model that has been developed previously and extensively applied to protein-DNA complex systems [20, 21, 22, 23, 24, 25, 26, 27, 28]. We used AICG2+ model for proteins, 3SPN.2 model for DNA [29, 30]. Briefly, in AICG2+, each amino acid in proteins are represented by one CG particle placed at its Cα position and the structure-based contact potential biases its energy landscape towards the reference structure. In 3SPN.2 model, each nucleotide is modeled by three CG particles corresponding to the phosphate, the sugar, and the base. Orientation-dependent potentials for base-base interactions and others are designed to reproduce basic experimentally-characterized properties of duplex and, to some extent, single strands. Between proteins and DNA, we applied the structure-based contact potential for representing the specific interactions, as well as a general excluded volume term and the electrostatic interaction via the Debye-Huckel approximation (the monovalent salt concentration was set to 200 mM throughout this study). For the electrostatic interaction, we employed partial charges on the surface residues of proteins, which were optimized to reproduce the electrostatic potential around the protein obtained by the all-atom model via the RESPAC method [52]. For time propagation, including the solvent effect implicitly, we employed a simple Langevin dynamics at the temperature 300K. For all the CG-MD simulations, we used the software CafeMol 3.2 [53].

The specific protein-DNA interaction, i.e., the structure-based contact potential is, as usual, expressed as

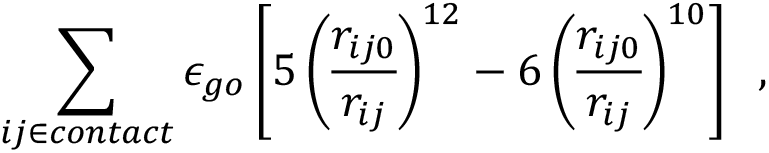

where *r*_*ij*_ is the distance between CG particles *i* and *j*, *r*_*ij*0_ is the corresponding distance at its reference structure, and *ϵ*_*go*_ is a parameter that modulates the strength of the interaction, of which value was calibrated to be 1.2 kcal/mol, to maintain experimentally characterized contacts in the three states, CC, OC, and ITC [10, 11]. Specifically, we set the following four conditions to be satisfied:

1. In the CC state, the contact between DNA and the E-wing of TFIIE is maintained.
2. In the OC state, the template DNA strand can maintain the native contacts with a region close to the active site of Pol II.
3. In the OC and ITC states, an upstream side of DNA maintains its contact with N50, K51, and T52 of TFIIE.
4. In the OC and ITC states, the triple mutations N50E, K51E, and T52E lead to loss of the DNA-TFIIE contacts [10].

### Trajectory analysis

The state-to-state transitions were characterized by protein-DNA contacts that depend on the state. There are 92, 22, and 118 contacts between proteins and DNA in CC, OC, and ITC states, respectively. The contacts in CC are all specific to the CC state and are not shared with the other two states. Thus, all the 92 contacts are used to characterize the CC state. The contacts in OC are mostly a subset of the contacts in the ITC state (19 out of 22 included in the ITC contacts). We used all the 22 contacts to characterize the OC state. Of the 118 contacts in ITC states, 99 are unique to the ITC state and thus are used to characterize the ITC state. Once the sets of contacts are defined, we quantify the state-to-state transition by the fraction of contacts formed in each snapshot.

The size of the DNA bubble was estimated as the sum of the broken base pairs, which are defined by the distance between the CG particles of the base pairs larger than 6.2 Å: In preliminary CG-MD simulations of duplex DNA at the same solvent condition, the probability that the base-base distance is larger than 6.2 Å was 0.3%. In apparently melted DNA configurations, their base-base distances were almost surely larger than this threshold distance.

### Back-mapping to all-atom model and all-atom MD simulations

Following the previously developed protocol [54], we performed the back-mapping from our CG models to all-atom models. While we used the cryo-EM-based CC protein structures as the reference structures of all the three states in the dynamical modeling, we moved them back to the respective cryo-EM protein structures aiming at more accurate modeling of all-atom structures. For the intermediate state I_2_, we used the OC structure as the reference. For each state, we began with the CG-MD simulation at 300 K for 10^5^ MD steps. Then, to reduce local fluctuations, we performed a quick annealing simulation, quenching the temperature from 300 K to 1 K, followed by a 10^5^ MD step simulation at 1 K. The final structure was put into the back-mapping toolset. For DNA, we applied the CG to AA reconstruction tool [54], whereas for proteins, we employed the PD2 ca2main [55] for backbone and SCWRL4 [56] for sidechain reconstruction.

Once the all-atom model for the PIC were obtained, we performed all-atom MD simulations using the software GROMACS 2020.2 [57] with the protein, DNA, and water force fields, ff14SB [58], and parmbsc1 [59], and TIP3P [60], respectively. We used the standard protocol: We set the box size of 182.2 × 232.1 × 186.4 Å^3^ solvating with water molecules and 171 Na^+^ ions to neutralize the system. After the energy minimization, we equilibrated the local system with NVT and then NPT ensembles (T= 300 [K], the pressure 1 bar), followed by 10 ns MD simulations. We used the cutoff distance of 1 nm for the Coulomb interaction with the particle-mesh-Ewald for long range treatment.

## Supporting information

Movie_S1_CCtoOC

Movie_S2_OCtoITC

---

**Figure S1:**
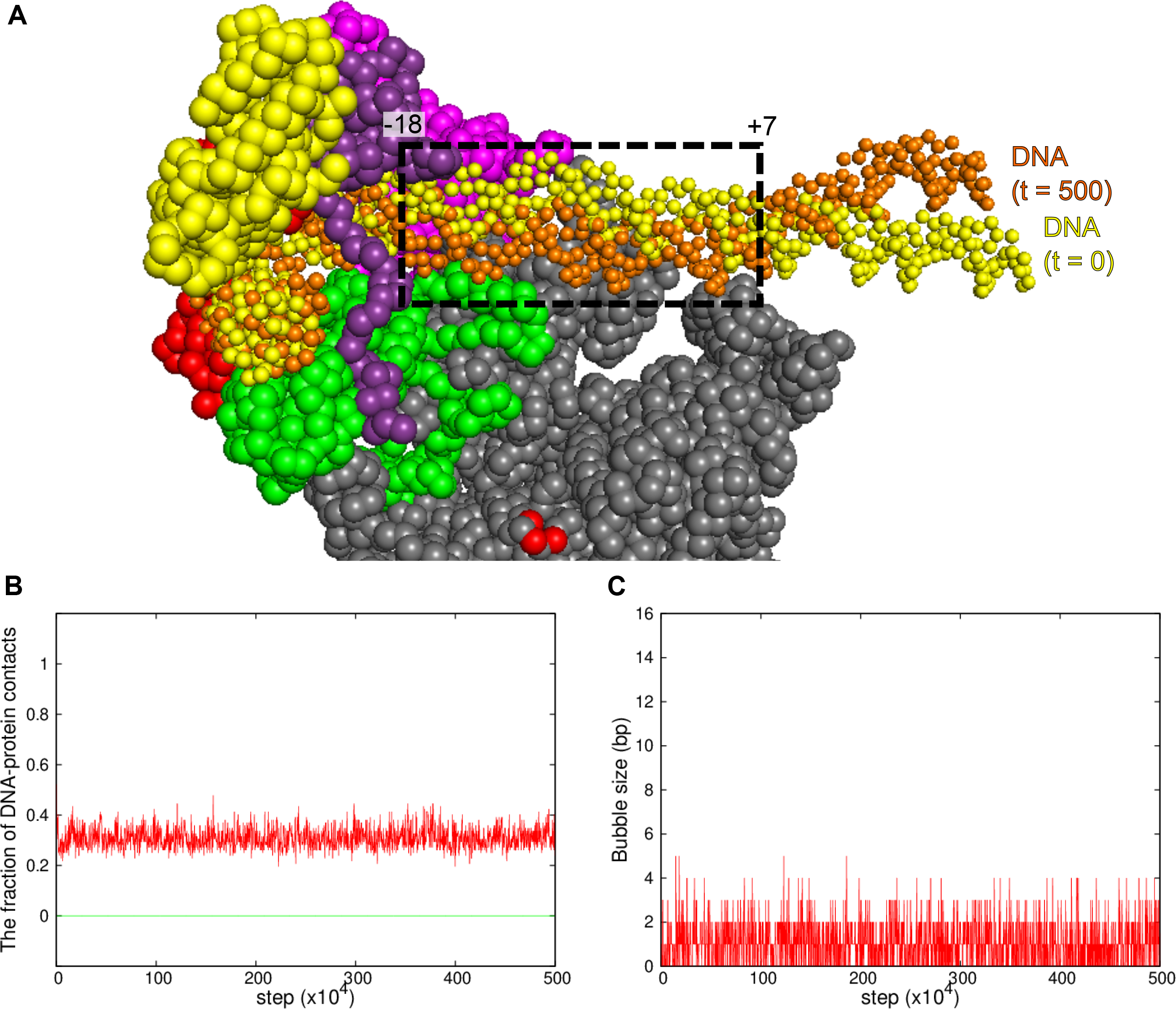
Coarse-grained MD simulation from the CC state without the mismatch DNA sequence. Results of a representative trajectory is shown. (A) Superposition of DNAs in the initial (t=0, yellow) and final (t=500 × 10^4^ MD steps, orange) states. Black dashed square indicates the distorted region of the promotor. (B) The time course of the fractions of protein-DNA contacts specific to CC (red) and OC (green). (C) The time course of the DNA bubble size.

**Figure S2:**
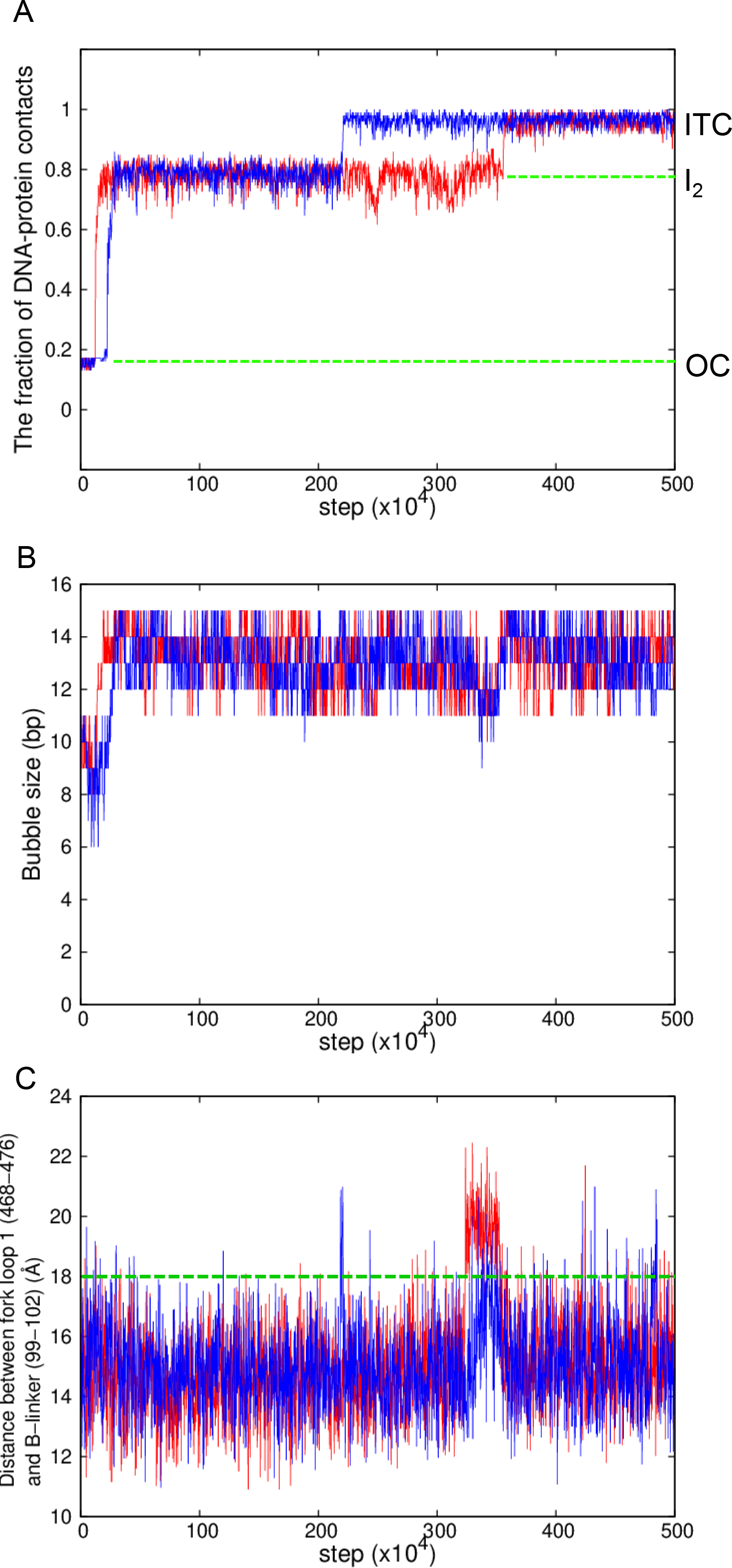
Coarse-grained MD simulation for the transition from the OC to ITC states with a weakened attraction between DNA and the E-wing residues. Results of typical two trajectories are shown in red and blue curves. (A) The time course of the fractions of protein-DNA contacts specific to ITC states. (B) The time course of the DNA bubble size. (C) The time courses of the distance between the centers of mass of the fork loop 1 of the Pol II Rpb2 (468-476 residues) and the B-linker in TFIIB (99-102 residues). Green dashed lines, a characteristic distance for the template DNA to pass through the fork loop 1.

**Figure S3:**
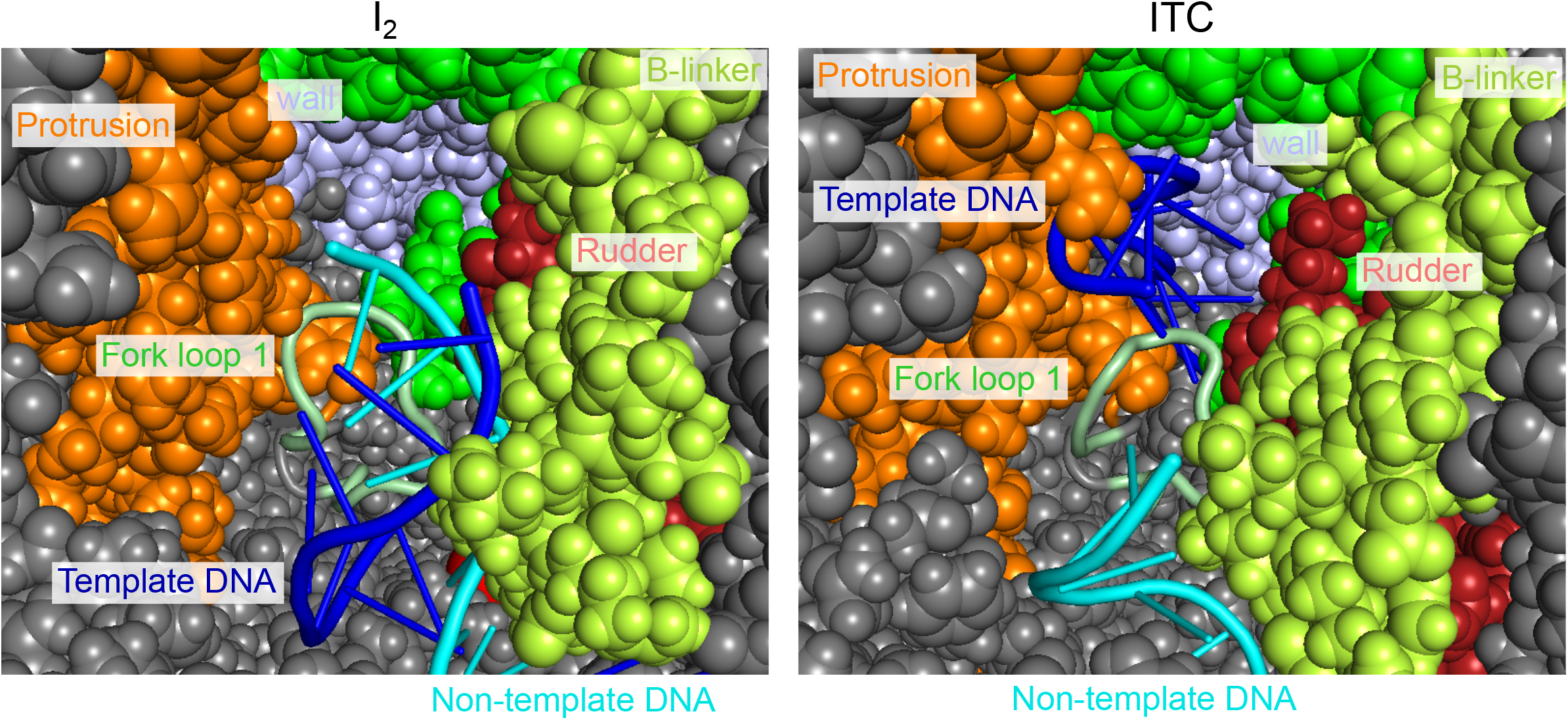
Close-up views around the fork loop 1 (pale green) of the Rpb2 in the I_2_ (left) and ITC (right) states. In the I_2_ state, both template and non-template DNA strands are in the near side of fork loop 1, whereas, in the ITC state, template and non-template DNA strands are in the near-side and the other side of the fork loop 1.

**Figure S4:**
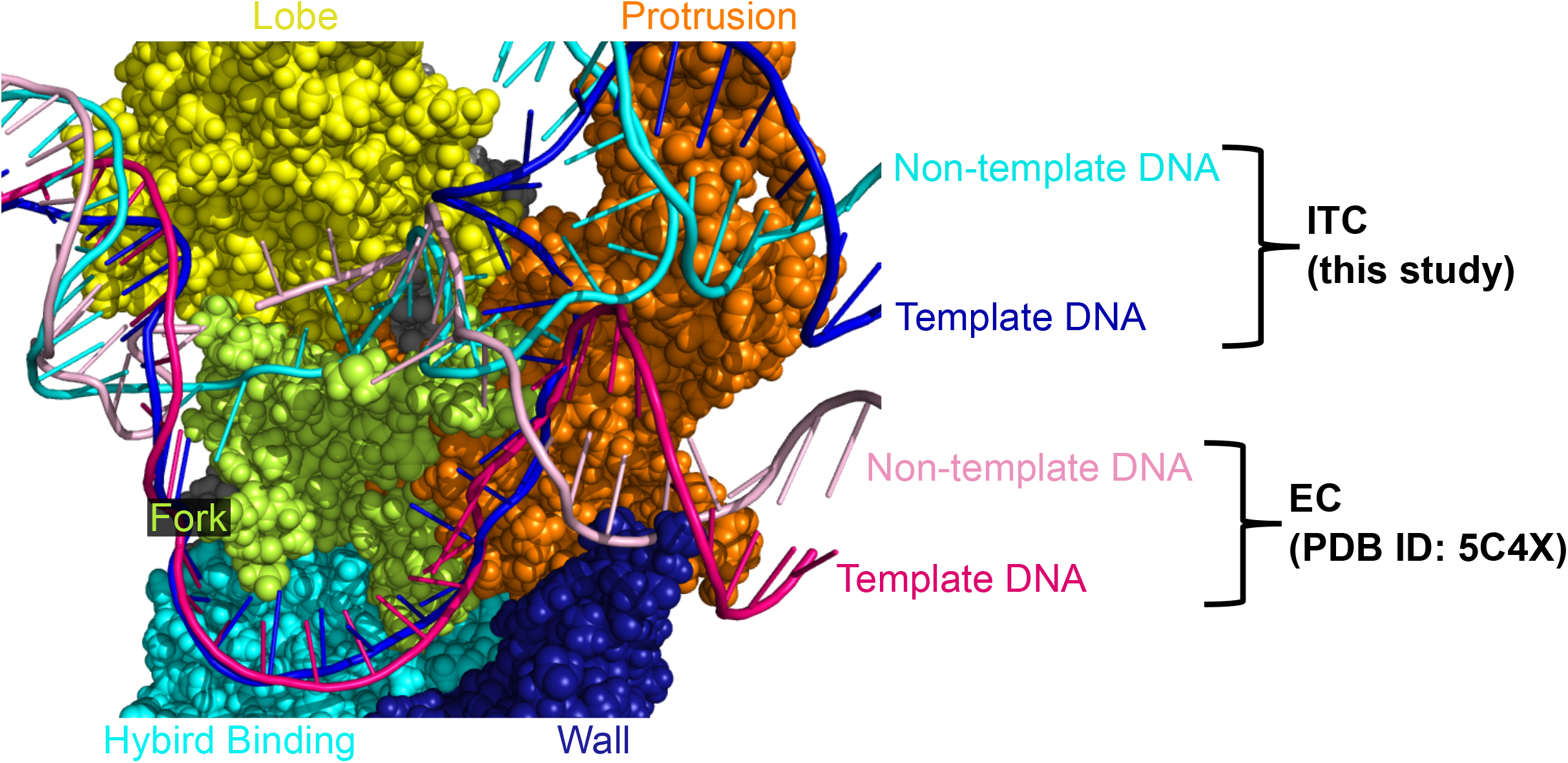
Comparison of the DNA bubbles in ITC (blue and cyan) and in the elongation complex (EC; PDB ID 5C4X) (magenta and pink). The DNA bubble structures in the two cases are similar.

**Figure S5:**
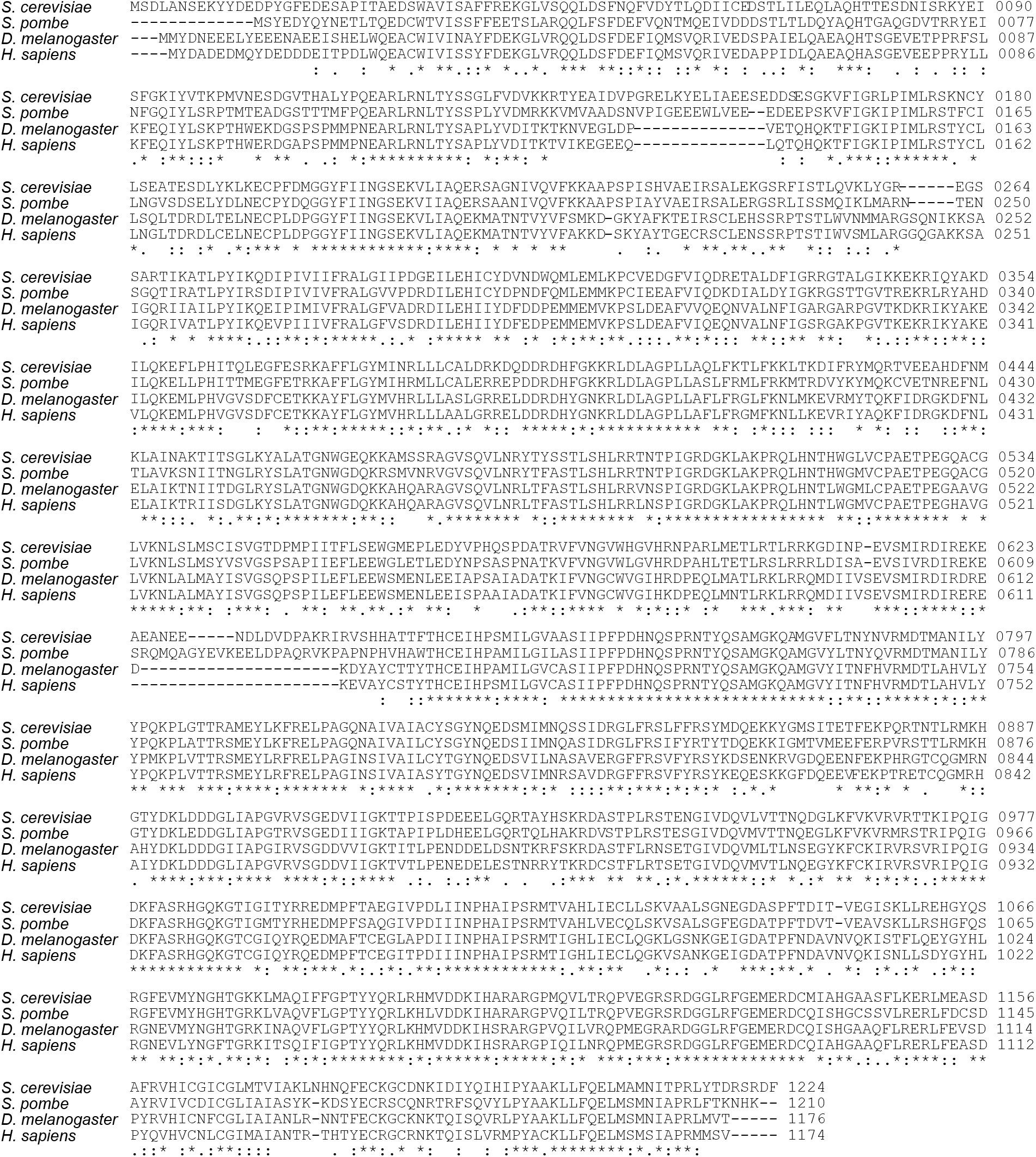
Multiple sequence alignment of Rpb2. “*”, identity; “:”, strong similarity; “.”, weak similarity.

**Movie S1: Coarse-grained MD simulation for the transition from the CC to OC states.**

**Movie S2: Coarse-grained MD simulation for the transition from the OC to ITC states.**

## Notes

### Competing Interest Statement

The authors have declared no competing interest.

